# Coral reef habitat complexity decreases and diversifies local availability of light

**DOI:** 10.1101/2024.12.12.628134

**Authors:** Viviana Brambilla, Joshua S. Madin, Nader Boutros, Luisa Fontoura, Mia Hoogenboom, Oscar Pizarro, Damaris Torres-Pulliza, Stefan Williams, Rachael Maree Woods, Kyle J. A. Zawada, Maria Dornelas

**Affiliations:** Centro de Ciências do Mar e do Ambiente, Faculdade de Ciências da Universidade de Lisboa, Lisboa, Portugal; Centre for Biological Diversity, University of St Andrews, St Andrews, UK; Hawai’ian Institute of Marine Biology, Kaneohe, HI, USA; Australian Institute of Marine Science, Perth, Australia; Macquarie University, Sydney, Australia; James Cook University, Townsville, Australia; Australian Centre for Field Robotics, University of Sydney, Sydney, Australia; Norwegian University of Science and Technology, Trondheim, Norway; Pacific Island Fisheries Science Center, NOAA Fisheries, Honolulu, HI, USA; NSW Department of Planning, Industry and Environment, Sydney, Australia

**Keywords:** Niche construction, structural complexity, geometric ecology, ecosystem engineers

## Abstract

The spatial distribution of environmental conditions can determine where organisms can and cannot live. When the distribution of microhabitats within a landscape is mediated by the shape of its surface, structural complexity can indirectly affect environmental niche distributions. This study investigates how light availability patterns change along a gradient of landscape complexity measured as surface rugosity and height range in tropical shallow reefs. We used 260,000 high frequency and high spatial resolution light measurements across 903 locations to determine the proportion of light that is available at the reef surface. We find that light available on the reef surface is highly variable, with close locations within reef sites experiencing up to 25.7 mol/m^2^d of daily photon input difference. After accounting for light attenuation due depth, light availability decreases with increasing surface rugosity: a unit increase in local surface rugosity corresponds to an 11-29% decrease in light availability. Local surface complexity can affect the distributions of light availability at local scales, while broader extent site metrics do not capture this variability. Our results suggest that structural complexity enhances local environmental variability, and its indirect effects on other environmental variables and their interactions are essential for understanding ecosystem processes.

## Introduction

Most life in the biosphere depends on solar radiation as a primary source of energy ^1^. With photosynthesis being the largest process of primary production, light availability is a key resource for most ecosystems, both marine and terrestrial. In surface ecosystems (terrestrial and benthic), ecosystem engineers create structures that mediate the distribution of this resource ^2,3^. How much light is available on the ecosystem surface is modulated by the structures created by the trees in forests, kelps in kelp forests, and corals on coral reefs. Here, we ask how the shape of these structures affects light availability on benthic substrata in coral reef ecosystems.

The energy stored through carbon fixation via photosynthesis uses only a fraction of the incoming solar radiation reducing the available energy for life ^1^. First, not all wavelengths are liable to be captured by the photosystems, with the fraction of light used for photosynthesis being known as the Photosynthetically Active Radiation wavelengths (PAR, 400-700 nanometer), a subset of the visible light. A common way to quantify light niches is then to measure the number of photosynthetically active photons (photons in the PAR frequency range) accumulated in an area over 24 hours (mol m^−2^ d^−1^), known as the daily light integral. Second, the atmosphere and water (in the case of aquatic ecosystems) absorb and scatter light ^4^. In oligotrophic marine water, irradiation scattering can be minimal because of the small particle and organismal densities of their waters. In most reefs, clear water bodies, where absorbance would mainly be due to water particles, the downward irradiance can be modelled as an exponential decrease with depth, with a coefficient of attenuation, *k*_*d*_, that depends on depth, wavelength range and water properties ^4^. Once *k*_*d*_ is known for a specific light wavelength, assuming a flat-water surface and clear water, light values should depend on depth and irradiance at the surface. Irradiance at the surface itself varies according to angle of incidence (time of day and seasonality), cloud cover and weather conditions in general. Third, the geometry of the ecosystem surface can also filter, intercept, or deflect light, and this process is the focus of this paper.

As on land, physical structures built by physical ecosystem engineers underwater give a third dimension to the landscape, potentially affecting local environmental conditions and increasing microhabitat diversity ^5^. Ecosystem surface elevations can disrupt light flow and cast shadows in the system. As most inhabited surfaces on Earth are not flat, the amount of light that can be available at any given location will be determined not only by elevation or depth, but also by the geometry of the local surface. A structurally complex site is likely to have more shadowed locations than a flat site, where depth is the main factor attenuating light that reaches the ground. With the possibility to digitalize benthic surfaces, complexity can now be defined through a set of continuous variables that capture different aspect of the geometric features of marine habitats ^6,7^. We can thus make concrete hypotheses for how light availability is affected by these geometric variables. For example, we can hypothesize that substrata with higher elevation range (height range) is more likely to produce shadowed environments. Surface rugosity [the ratio between the 3D surface the reef and its planar projection on the horizontal plane ^8^] can also reduce the amount of available light at location.

Coral reefs harbour some of the most biodiverse and threatened communities in the world ^9,10^. Although corals are animals, they use light as their main source of energy. Most reef-building corals rely on an intimate symbiosis with dinoflagellate algae, called zooxanthellae, for energy supply and need the right environmental conditions to have an efficient symbiosis ^11^. Like their plant counterparts, light and temperature are two particularly important environmental variables for photosynthesis outputs. In fact, zooxanthellate corals rely on photosynthesis outputs for up to the 95% of their energetic input and photosynthesis is tightly linked to calcification ^12^. However, if local light and temperature increase above species physiological thresholds, the zooxanthellae get expelled from the coral causing bleaching, a starving condition that can lead to coral mortality ^13^. Temperature and solar radiation thresholds for coral bleaching have been experimentally defined for several species and vary across taxa ^14^. High seawater temperatures have been shown to increase bleached coral cover in the field ^15^. However, the cause for small-scale spatial patchiness in bleaching on reefs remains unclear and could be linked to small scale habitat variation. Light availability is a primary factor of influence of coral growth and calcification, as coral in higher light environments on average grow three times more than in darkness ^12^. Furthermore, light plays an important role in determining colony morphologies and resource allocation ^16^. For example, foliose corals respond to light gradients maximizing the planar area of the colony ^17^ and massive corals change the geometry of corallites in order to harvest more light ^18,19^.

Coral reef structures and functioning rely on the production of calcium carbonate through skeleton accretion of the hard coral inhabiting them, which form the basis of the reef matrix. Corals in reef ecosystems are structural obligate ecosystem engineers and modify the ecosystem by their own physical structure without the possibility to do otherwise ^2^. As such, optimal environmental conditions for coral performance in reefs are of primary importance to enable reefs to persist and function. At the same time, local environmental conditions are likely to be mediated by the coral structures themselves as the distribution of microhabitats within a reef landscape can be mediated by the shape of its surface. Smooth flat surfaces are expected to have more homogenous environments, whereas convoluted, complex surfaces have more variable environments. In this study, we look at light availability patterns along a gradient of landscape complexity in tropical shallow reefs. We predict that the proportion of light available on the reef substratum decreases with complexity, measured as surface rugosity and height range at different local scales. Finally, we predict that the relationship between surface geometry and light availability is scale dependent, with local scale complexity having a stronger effect on light availability, than coarser grain site level metrics.

## Methods

### Study sites

Data were collected at nine shallow reef flat sites encircling Jiigurru (Great Barrier Reef, Australia, Figure 1a) in November 2017, 2018, and 2019. Sites were about 130 m^2^ in area and were chosen to capture a range of habitat and structural conditions (Torres-Pulliza et al. 2020). Digital elevation models (DEMs) and orthomosaics were created for each site using structure from motion photogrammetry (Figure 1c). We used the spiral method described in Pizarro et al. ^20^ and the processing pipeline described in Torres-Pulliza et al. ^7^.

**Figure 1.**
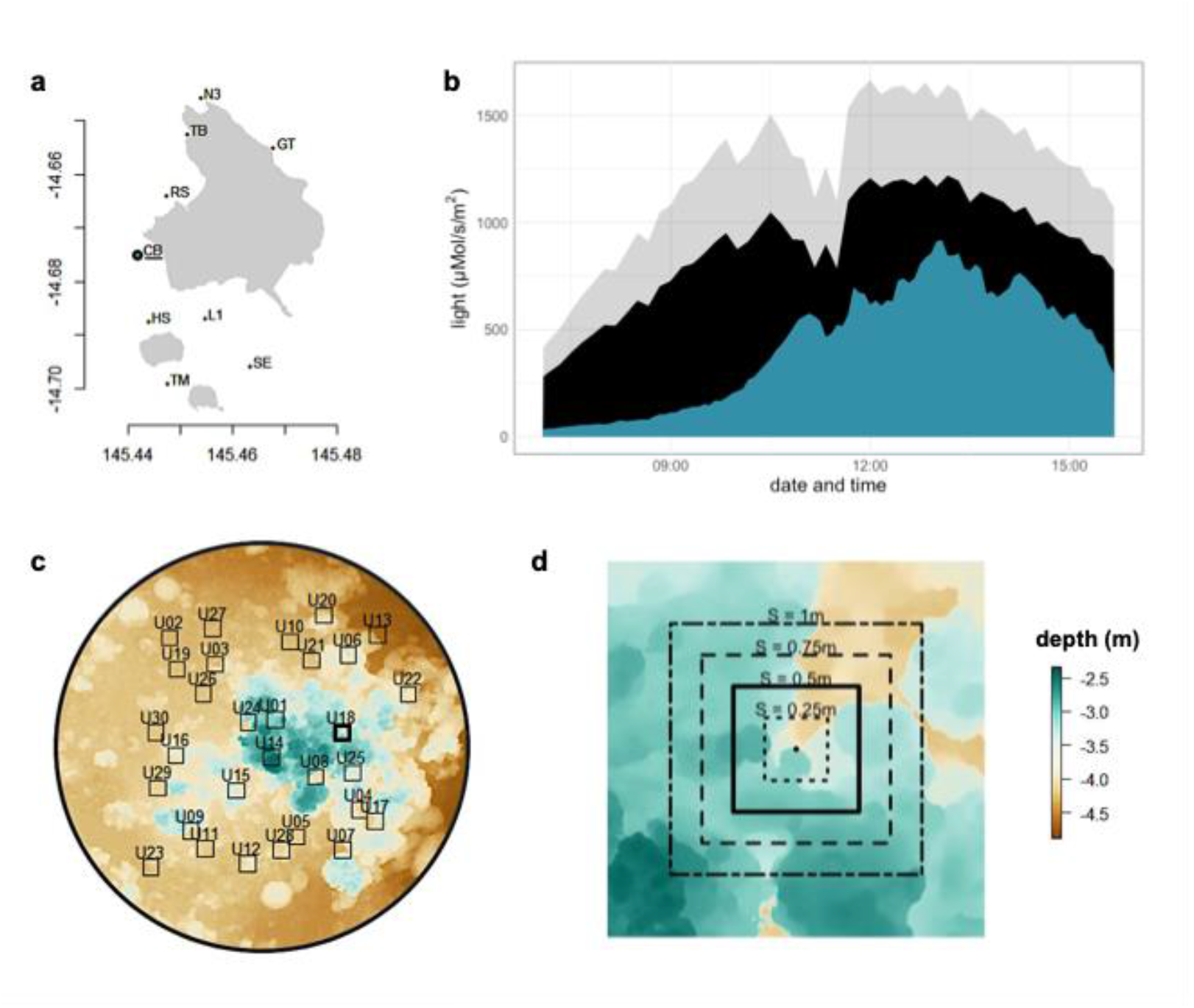
Study sites and sampling scheme example. **(a)** Map of sampling sites (CB= Corner Beach, GT = Gnarly Tree, HS = Horseshoe, L1 = Lagoon 1, N3 = North Reef 3, RS = Resort, SE= Southeast, TB = Turtle Beach, TM = Trimodal) around Jiigurru (Australia). **(b)** Example light distributions during a sampling event for one sampling point. In grey, the photosynthetically active radiation (PAR) measured by AIMS meteorological buoy. In black, the PAR expected at depth of sensor deployment, attenuated following eq.1. In blue, the PAR as measured *in situ* with loggers at substratum. The response variable is the proportion between the PAR as measured *in situ* with loggers at substratum (blue area) and the bottom attenuated PAR (black area). **(c)** A digital elevation model for Corner Beach, showing an example of the randomised locations within site where PAR data loggers were deployed during a sampling event. Squares have sides of 50 cm and are centred on the location where the logger unit was deployed. **(d)** Local scale sizes considered in the analysis. The central dot represents the position of the logger and the concentric squares representing the different local scales considered in extraction of surface geometry metrics.

### Light

Every year, five to nine sampling events on different days, one per site, were performed and light was recorded at 30-50 randomly selected surface locations within each site and year using HOBO Onset pendant data loggers. The data loggers were attached to dive weights and deployed haphazardly within the sites with the light sensor facing directly upwards and horizontally orientated (Figure 1c). The positions of loggers were annotated on printouts of site orthomosaics. If loggers changed position or inclination during the sampling period, they were excluded from the analysis. Loggers recorded light (lux) values every five minutes for 24 hours (i.e., the sampling event) and we converted these values into PAR (in μMol/m2/s) using the conversion coefficient in Thimijan and Heins (1983) ^21^ This conversion is unreliable for absolute values of PAR, but relative values are robust. For each sampling event and logger, we computed the recorded daily light integral.

To measure daily integral of PAR expected at sampling depth at each sampling event, we used the downward expected irradiance built with an attenuation function (Kirk 1983) that computed an exponential decrease of light given light available at surface and depth of sampling:

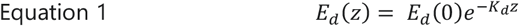

Where *E*_*d*_ is the downward irradiance, *z* is depth, and *k*_*d*_ is the coefficient of light attenuation, which we assume constant. To compute depth at each sampling timestep at each location, water column height above each logger location was obtained by combining depth of DEM and the current tidal phase. Tide prediction at each round hour and maximum and minimum tide time were available from the Queensland Government website ^22^. To predict continuous tidal regime for the whole sampling event duration, the ‘TideHarmonics’ package ^23^ was used, and the available data was interpolated with 37 harmonics, as suggested for mixed semidiurnal tide regimes (i.e. the tidal regime of Jiigurru). Surface PAR, *E*_*d*_*(0)*, was collected at an elevated weather station and available online at the Australian Institute of Marine Science Data Centre ^24^. Thus, at each timestep the available PAR per depth was computed and the daily integral of PAR expected at each sampling location was computed as the daily integral (Figure 1b).

The proportion of the recorded daily light integral with respect to the expected daily light integral for that sampling event day was used as response variable.

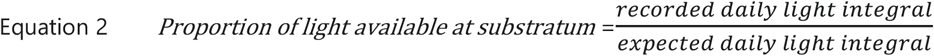

### Habitat complexity

Geometric traits of the reef, this is surface rugosity and height range, were calculated for square patches centred on each data logger for four different sizes S: 0.25 m, 0.5 m, 0.75m and 1m (Figure 1d). The 0.5 m patch was chosen to represent a medium benthic sessile organism size, and the smaller and larger patches were chosen to test for scale dependence. Surface rugosity was calculated as the ratio between the 3D area of the surface of each cropped patch obtained with the ‘surfaceArea’ function in the package ‘sp’ ^25^ and its planar area (i.e., S^2^). Height range was calculated as the difference between the lowest and highest elevation point within each patch. Site level traits were also computed for the central 8m × 8m square centred in each site map central coordinates by averaging four values obtained in their 4mx4m quadrants ^7^.

All the analysis of the DEMs were carried out with R, version 3.6.0 (R Core Team, 2018), using the raster and sf packages ^26,27^, where not specified otherwise.

### Analysis

Bayesian logistic mixed models were fitted to examine the effect of surface rugosity and height range on explaining the proportion of light available at substratum. A model per each size of the area considered to compute complexity (S = 0.25 m, S = 0.5 m, S = 0.75 m, S = 1m and S = 8m, i.e. site level) was fitted. Because of the non-independence between surface rugosity and height range ^7^, the interaction effect between those variables was included in the model. Sampling event was included as random effect to account for unexplained differences across sampling conditions.

Models were fitted with the probabilistic language RStan using the ‘brms’ package ^28^. We specify a beta family to account for the distribution of the response variable (proportion,^29^). All the priors were left as default, and for each model four chains for 4000 iterations were run, with a warm-up period of 1000 iterations. Convergence and stability of the Bayesian sampling was assessed using R-hat, which should be below 1.01 ^30^, and Effective Sample Size (ESS), which should be greater than 1000 ^28^. All analyses were performed in R version 3.3.2 ^31^.

## Results

260,000 light measurements were taken across 903 reef locations and over 23 sampling events over the course of this study (Figure 1, Figure 2). The proportion of light available at each location was obtained after computing light daily integrals of the *in situ* light measurements and light attenuation curves based on time of recording and depth of each individual location (Figure 1). Overall, there was substantial within site variation in proportion of light available and in most cases, some locations had almost no light for the whole sampling event (Figure 2). Across sampling events, local daily integrals differed in daily photon input by 9.05 to 25.69 mol/m^2^, which corresponded to ranges of 35.5 to 86.7% of the possible available light at depth. With regard to structural complexity, local estimates of height range and surface rugosity covered gradients spanning up to three orders of magnitude at each scale considered (Figure 1, Figure 2).

**Figure 2.**
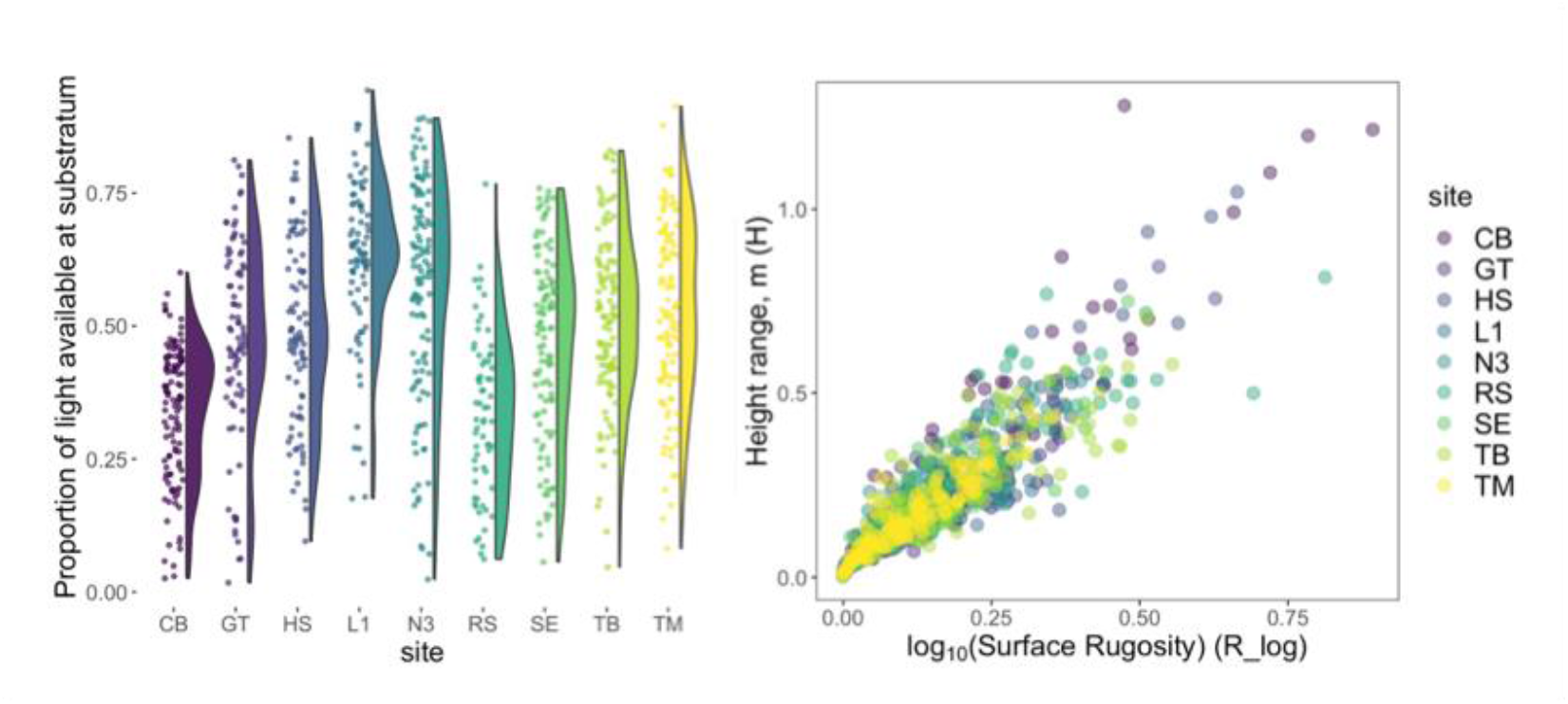
Available light and complexity variables distributions. On the left, raincloud plot of the proportion of light available at substratum, with sites ordered by increasing average surface rugosity. On the right, structural complexity variables distributions for height range (H) and surface rugosity (R, logged) referring to the area of a square of 50 cm of substratum centred on the position of the logger. Sites are ordered by site level structural complexity. (CB= Corner Beach, GT = Gnarly Tree, HS = Horseshoe, L1 = Lagoon 1, N3 = North Reef 3, RS = Resort, SE= Southeast, TB = Turtle Beach, TM = Trimodal).

The model for each size considered converged (R-hat = 1.000, all models). We found a consistent negative effect of surface rugosity at smaller local scales (S=0.25m, S=0.5m, S=0.75m, Figure 3). We could not detect an effect of the interaction between height range and surface rugosity at any scale considered, nor an effect of height range on estimated proportion of light available at substratum (Figure 3).

**Figure 3.**
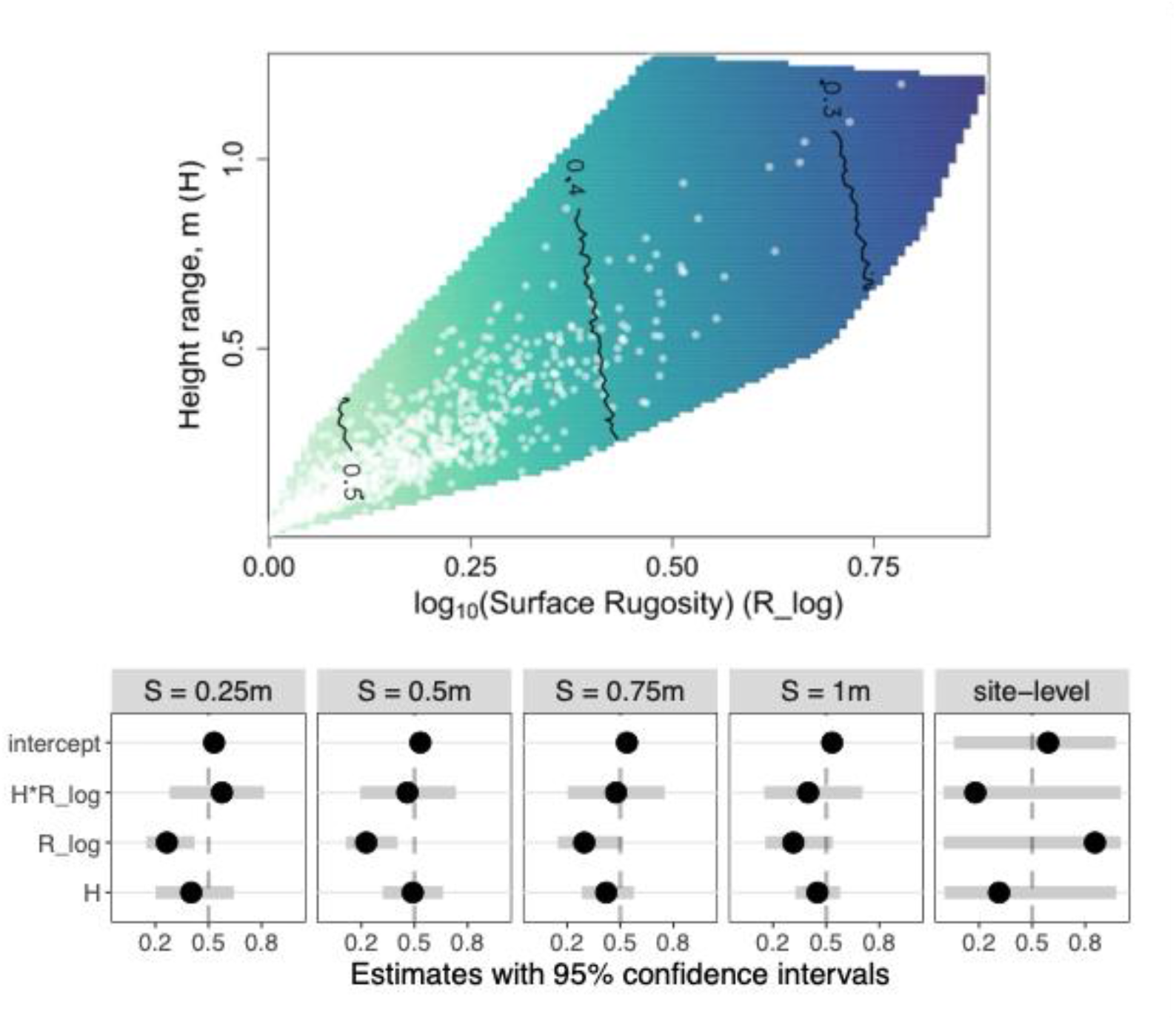
Proportion of available light distributions across the structural complexity space measured for areas of 0.5m and models results across scales. Contour plot of the effects of structural complexity on the proportion of light available at S=0.5m. Lighter areas represent complexity configurations with higher predicted proportional light available at substratum. On the bottom panel, the dots represent model effect sizes, and the grey bars represent 95% credible intervals per each size of area considered. Notice that values have been back-transformed to be more informative, so the dotted line lies at 0.5.

In the model fitted at the 0.5m scale (Figure 3), the model’s explanatory power was substantial (R^2^= 0.35, 95% CI = [0.31, 0.39]). Within this model, rugosity had 99.79% probability of having a negative effect based on posterior distributions, with a median posterior distribution of -1.23 (95% CI = [-2.09, -0.39]), meaning that a unit increase in surface rugosity corresponds to a decrease in light availability of 23%. The interaction effect was overall negative but had a very large credible interval (median posterior distribution = -0.15, 95% CI = [-1.43, 1.07]), which resulted in almost a half split between the effect being overall negative or positive (59.24% probability of being negative). The effect of height range was also overall negative, but almost centred in 0 (median posterior distribution = -0.03, 95% CI = [-0.76, 0.68]), meaning that we did not detect a reliable directional effect of this variable.

Across the other local scales (S=0.25m, S=0.75m, S= 1m), the median of the posterior distribution of the effect of rugosity ranged from -0.88 to -0.78 (Figure 3, SM1). When fitting site level complexity variables, surface rugosity had a very large credible interval (median posterior distribution = 1.72, 95% CI = [-9.16, 12.80]) and we did not detect a directional effect of this variable, or any variable at all (SM1, Figure 3).

## Discussion

In this study we quantified the effect of reef complexity on light availability using high frequency and high-resolution *in situ* light measurements across shallow reef sites. Light availability was highly variable within sites, and declined on average with increasing complexity, namely surface rugosity, at local scales. We found high variability in light availability within sites during each sampling event: most sites experience almost full shadow at least in some sampled locations throughout the day, while also having some locations experiencing close to the maximum available light they can experience at their given depth. Our findings align with previous research on the role of structural complexity in shaping environmental conditions ^20^ and ecological niches. Effects of local complexity within reef sites on light availability alters environmental regimes and niches variability at local scales, which can have cascading effects on benthic communities, their dynamics and primarily their primary production ^21^.

Our results highlight the potential for structural complexity to mediate ecosystem responses to environmental change by modulating light availability. Light can have substantial effects on benthic metabolisms and primary production ^17,22–24^. Specific to this system, less extreme conditions in light regimes are typically optimal for most corals’ metabolisms and growth ^14,16^ and our results suggest presence of a range of these conditions is maximized in locally complex reefs. Additionally, the high variability in light availability and the local reef complexity might also explain the high coral morphological diversity found in small extents of reefs ^16,18,25^.

In addition to its effect on light availability, structural complexity affects other environmental variables and reef ecosystem processes. For example, higher rugosity increases the surface available for colonization. Complexity affects nutrient distribution and heat fluxes ^26^ through its effects on flow regimes. High surface rugosity has been associated with eddies and turbulent conditions ^20,26–28^. In combination, these relationships suggest structural complexity may help explain patchiness in bleaching.

In this study we measured complexity with height range and surface rugosity across scales, but these two variables are not completely independent ^7^. The lack of independence explains why when surface rugosity was removed from the models, we found similar effects of height range on light availability at substratum across scales (SM2). Results were not only robust to the choice of variable chosen to describe complexity but for each model fitted, they maintained their scale dependence as well (SM2). As spatial scale increased, the credible intervals around effect sizes increased, decreasing our ability to find a signal in this relationship. Nonetheless, results at scales comparable with an average adult coral colony size (S=0.50m and S=0.75cm) were largely consistent (Figure 3, SM1, SM2). The presence of an adult colony and the coral community structure itself can then play a huge role on the distribution of light availability within reefs. This has implications for the design and monitoring of applied science (restoration) and ecosystem functioning studies. For instance, the resolution of the maps used in this analysis is orders of magnitude higher than what many large scale boat-mounted benthic monitoring projects produce, which can overlook those fine details^29^. To protect and manage biodiversity and ecosystems, conservation efforts, especially local ones, should include finer scale structural complexity monitoring approaches, as using coarser measures might overlook important feedback loops that might occur at a smaller scale^29^. If we are to understand ecosystem functioning, we need to measure variables at the appropriate scale and consider the dual relationship between organisms and the environment.

Despite the clear role of surface rugosity, most of the variation in light availability remains unexplained. Future work should explore the role of turbidity, and water surface smoothness on light availability, all of which have been shown to influence light availability in benthic substrata ^30,31^. Specifically, turbidity should also be considered when computing the light attenuation curve as it can affect the k_d_ parameter that defines attenuation ^4^. Turbidity has usually greater effects with increasing depths ^30^ and could explain why light proportions are skewed towards lower values in sites that are overall deeper (Figure 3). In fact, the deepest sites monitored (Corner Beach and Resort) also correspond to sites that have higher proportion of sand substratum and longer distance to fully consolidated reef, which also contributes to regulate particle suspension, and in turn control clearness of the water ^4^.

The spatial distribution of environmental conditions can determine where organisms can and cannot live ^32^. Mechanistically, environmental conditions can regulate local species occurrence and performance by influencing biological and ecological processes such as physiological status ^33^, recruitment ^34^, competition ^35^, and predation ^36^. Here we show that the distribution of light microhabitats within a landscape is mediated by the shape of its surface, creating direct effects or buffers with respect to overall environmental conditions. Like on land, where the makeup of trees sizes and species determines light heterogeneity and drives forest succession ^37^, benthic organisms have the potential to start cascading effects on the local community by modifying the surrounding environment. In shallow benthic habitats such as coral reefs or kelp forests, this is particularly significant as the shape of the landscape is often built by organisms whose shape features can determine light partitioning across the surface. Corals and the reefs they leave behind intercept and change something as fundamental to life as light, and these changes contribute to the diversity of environmental conditions found on the reef. Foundational species, and ecosystem engineers in general, are thus instrumental agents in constructing their own niche.

## Acknowledgments

We thank the Australian Museum’s Lizard Island Research Station staff and S.Patzy for their support. VB would like to thank members of the Fish Lunch discussions along the years and the writing retreat organized at the Hawai’ian Institute of Marine Biology in 2022. Funding was provided by the Warman Foundation (to MD and JSM), the John Templeton Foundation (MD, JSM, MH, grant #60501 ‘Putting the Extended Evolutionary Synthesis to the Test’), a Royal Society research grant and a Leverhulme fellowship (MD), two Ian Potter Doctoral Fellowships at Lizard Island Research Station (DTP, VB) and a MASTS small grant (VB). Partly funded by the European Union (CoralINT, GA 101044975). Views and opinions expressed are however those of the author(s) only and do not necessarily reflect those of the European Union or the European Research Council. Neither the European Union nor the granting authority can be held responsible for them.

## Supplementary material

**SM1.**
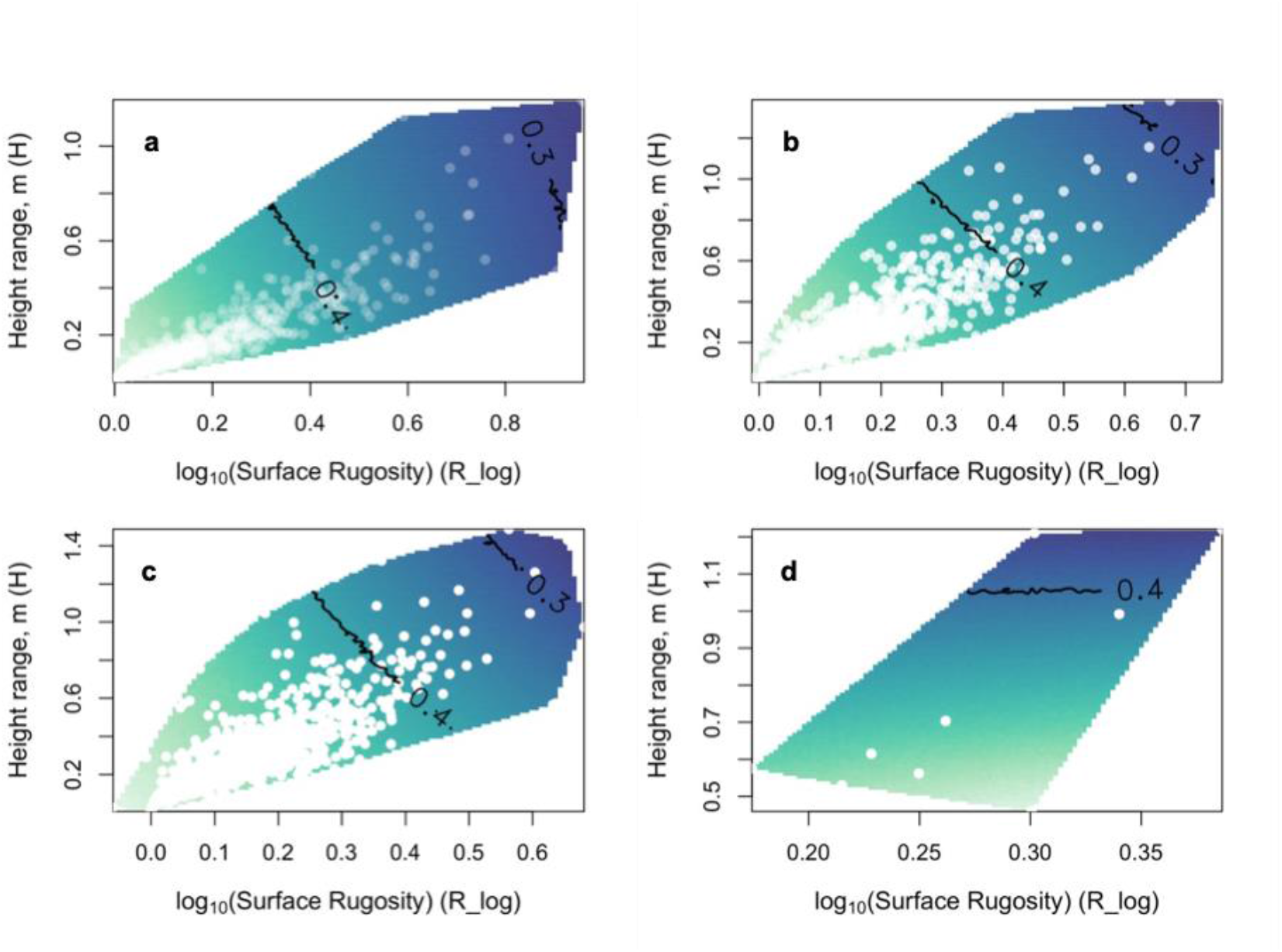
Predictions of light availability at substratum for all models fitted. a) S=0.25m, b) S=0.75m, c) S=1m, d) site level metrics.

**SM2.**
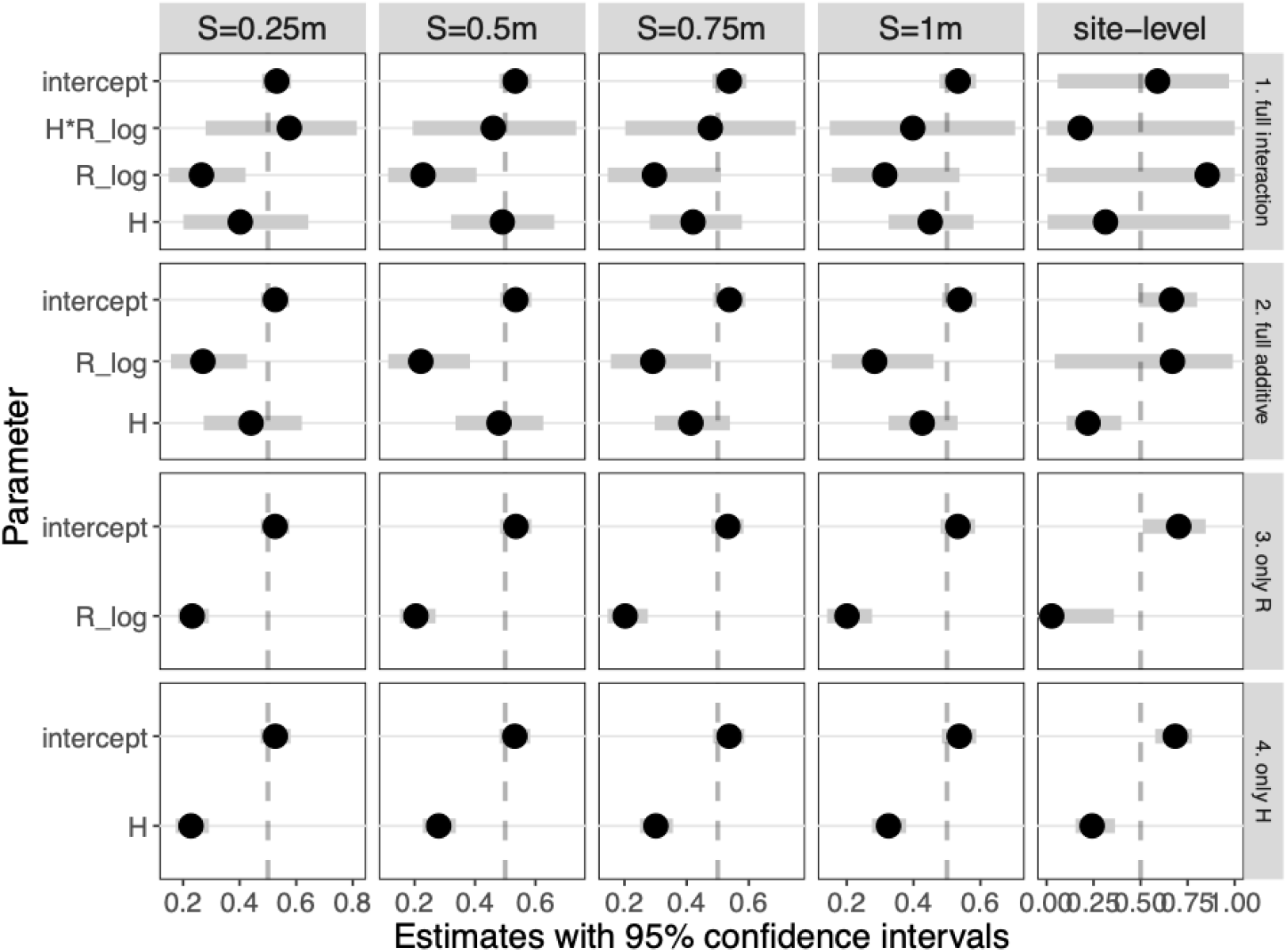
Sensitivity analysis and effect sizes for all models fitted. Nested hierarchical models were fitted and leave-one-out cross validation used to assess predictive performance of the models. Differences in expected log predictive density (ELPD) were smaller than 4 for each scale considered, so the models have very similar predictive performance.

## References

1. Vernadskij, V. I., McMenamin, M. A., Langmuir, D. B. & Vernadskij, V. I. The Biosphere. (Copernicus, New York Heidelberg, 1998).

2. Jones, C. G., Lawton, J. H. & Shachak, M. Organisms as ecosystem engineers. Oikos 69, 373–386 (1994).

3. Jones, C. G., Lawton, J. H. & Shachak, M. Positive and Negative Effects of Organisms As Physical Ecosystem Engineers. Ecology 78, 1946–1957 (1997).

4. Kirk, J. T. Light and Photosynthesis in Aquatic Ecosystems. 401 (Cambridge University Press, Cambridge, UK, 1983).

5. Berke, S. K. Functional groups of ecosystem engineers: A proposed classification with comments on current issues. Integrative and Comparative Biology 50, 147–157 (2010).

6. Burns, Delparte, Gates, & Takabayashi. Utilizing underwater three-dimensional modeling to enhance ecological and biological studies of coral reefs. International Archives of the Photogrammetry, Remote Sensing and Spatial Information Sciences - ISPRS Archives XL-5/W5, 61–66 (2015).

7. Torres-Pulliza, D. et al. A geometric basis for surface habitat complexity and biodiversity. Nature Ecology & Evolution (2020) doi:10.1038/s41559-020-1281-8.

8. Friedman, A., Pizarro, O., Williams, S. B. & Johnson-Roberson, M. Multi-Scale Measures of Rugosity, Slope and Aspect from Benthic Stereo Image Reconstructions. PLoS ONE 7, (2012).

9. Veron, J. Corals in Space and Time: The Biogeography and Evolution of the Scleractinia. (UNSW Press, Sydney, Australia, 1995).

10. Hughes, T. P. et al. Coral reefs in the Anthropocene. Nature 546, 82–90 (2017).

11. Muscatine, L. Nutrition of corals. in Biology and Geology of Coral Reefs (eds. Jones, O. A. & Endean, R.) 77–115 (Academic Press, New York, USA, 1973).

12. Gattuso, J. P., Allemand, D. & Frankignoulle, M. Photosynthesis and calcification at cellular, organismal and community levels in coral reefs: A review on interactions and control by carbonate chemistry. American Zoologist 39, 160–183 (1999).

13. Brown, B. E. Coral bleaching: Causes and consequences. Coral Reefs 16, 129–138 (1997).

14. van Oppen, M. J. H. & Lough, J. M. Coral Bleaching: Patterns, Processes, Causes and Consequences. Chapter 1, 2 and 3 (2009). doi:10.1007/978-3-540-69775-6.

15. Hoegh-Guldberg, O. Climate change, coral bleaching and the future of the world’s coral reefs. Marine and Freshwater Research 50, 839–866 (1999).

16. Todd, P. A. Morphological plasticity in scleractinian corals. Biological Reviews 83, 315–337 (2008).

17. Hoogenboom, M., Connolly, S. & Anthony, K. Interactions between morphological and physiological plasticity optimize energy acquisition in corals. Ecology 1144-1154-1144–1154 (2008).

18. Bruno, J. F. & Edmunds, P. J. Metabolic consequences of phenotypic plasticity in the coral Madracis mirabilis (Duchassaing and Michelotti): The effect of morphology and water flow on aggregate respiration. Journal of Experimental Marine Biology and Ecology 229, 187–195 (1998).

19. Todd, P. A., Ladle, R. J., Lewin-Koh, N. J. I. & Chou, L. M. Genotype x environment interactions in transplanted clones of the massive corals Favia speciosa and Diploastrea heliopora. Marine Ecology Progress Series 271, 167–182 (2004).

20. Pizarro, O., Friedman, A., Bryson, M., Williams, S. B. & Madin, J. A simple, fast, and repeatable survey method for underwater visual 3D benthic mapping and monitoring. Ecology and Evolution 7, 1770–1782 (2017).

21. Thimijan, R. & Heins, R. Photometric, radiometric, and quantum light units of measure: a review of procedures for interconversion. Horticultural Science 18, 818–822 (1983).

22. Maritime Safety Queensland & Department of Transport and Main Roads. 2020 Tide Predictions Blue Book - Far North Queensland. 33–33 https://www.msq.qld.gov.au/Tides/Supplementary-tide-predictions.aspx (2019).

23. Stephenson, A. A. & Stephenson, M. A. Package ‘ TideHarmonics ‘. (2017).

24. AIMS. Data Centre. 2020.

25. Pebesma, E. & Bivand, R. Classes and methods for spatial data in R. R News 5, 9–13 (2005).

26. Hijmans, R. J. raster: Geographic Data Analysis and Modeling. R package version 3. 3–13. (2020).

27. Evans, J. S. Package ‘ spatialEco ‘. (2020).

28. Bürkner, P.-C. brms : An R Package for Bayesian Multilevel Models Using Stan. Journal of Statistical Software 80, (2017).

29. Douma, J. C. & Weedon, J. T. Analysing continuous proportions in ecology and evolution: A practical introduction to beta and Dirichlet regression. Methods Ecol Evol 10, 1412–1430 (2019).

30. Vehtari, A., Gelman, A.,. and Gabry, J. loo: Efficient leave-one-out cross-validation and WAIC for Bayesian models. (2016).

31. R Core Team. R: A language and environment for statistical computing. R Foundation for Statistical Computing (2023).

32. Hench, J. L. & Rosman, J. H. Observations of spatial flow patterns at the coral colony scale on a shallow reef flat. Journal of Geophysical Research: Oceans 118, 1142–1156 (2013).

33. Denny, M. Ecological Mechanics: Principles of Life’s Physical Interactions. (Princeton University Press, 2015). doi:10.2307/j.ctvc775jj.

34. Hoogenboom, M. O. & Connolly, S. R. Defining fundamental niche dimensions of corals : synergistic effects of colony size, light, and flow. Ecology 90, 767–780 (2009).

35. Lenihan, H. S. Physical-Biological Coupling on Oyster Reefs: How Habitat Structure Influences Individual Performance. Ecological Monographs 69, 251–251 (1999).

36. Luhtala, H., Kulha, N., Tolvanen, H. & Kalliola, R. The effect of underwater light availability dynamics on benthic macrophyte communities in a Baltic Sea archipelago coast. Hydrobiologia 776, 277–291 (2016).

37. Smith, L. W., Barshis, D. & Birkeland, C. Phenotypic plasticity for skeletal growth, density and calcification of Porites lobata in response to habitat type. Coral Reefs 26, 559–567 (2007).

38. Kaandorp, J. A. et al. Simulation and analysis of flow patterns around the scleractinian coral Madracis mirabilis (Duchassaing and Michelotti). 1551–1557 (2003) doi:10.1098/rstb.2003.1339.

39. Lowe, R. J., Shavit, U., Falter, J. L., Koseff, J. R. & Monismith, S. G. Modeling flow in coral communities with and without waves: A synthesis of porous media and canopy flow approaches. Limnology and Oceanography 53, 2668–2680 (2008).

40. Stocking, J. B., Laforsch, C., Sigl, R., Reidenbach, M. A. & Stocking, J. B. The role of turbulent hydrodynamics and surface morphology on heat and mass transfer in corals. Journal of the Royal Society Interface 20180448, (2018).

41. Lutzenkirchen, L. L., Duce, S. J. & Bellwood, D. R. Exploring benthic habitat assessments on coral reefs: a comparison of direct field measurements versus remote sensing. Coral Reefs 43, 265–280 (2024).

42. Anthony, K. R. N., Ridd, P. V., Orpin, A. R., Larcombe, P. & Lough, J. Temporal variation of light availability in coastal benthic habitats: Effects of clouds, turbidity, and tides. Limnology & Oceanography 49, 2201–2211 (2004).

43. Storlazzi, C. D., Norris, B. K. & Rosenberger, K. J. The influence of grain size, grain color, and suspended-sediment concentration on light attenuation: Why fine-grained terrestrial sediment is bad for coral reef ecosystems. Coral Reefs 34, 967–975 (2015).

44. Hutchinson, G. E. Concluding Remarks. Cold Spring Harbor Symposia on Quantitative Biology 22, 415–427 (1957).

45. Bradford, K. J. & Hsiao, T. C. Physiological Plant Ecology II. 324 (Springer Berlin Heidelberg, Berlin, Heidelberg, 1982). doi:10.1007/978-3-642-68150-9.

46. Underwood, A. J. & Denley, E. J. Paradigms, Explanations, and Generalizations in Models for the Structure of Intertidal Communities on Rocky Shores. in Ecological Communities, Conceptual Issues and the Evidence (eds. Strong, D., Simberloff, D., Abele, L. & Thistle, A.) 151–180 (Princeton University Press, Princeton, 1984). doi:10.1515/9781400857081.151.

47. MacArthur, R. H. & MacArthur, J. W. On Bird Species Diversity. Ecology 42, 594–598 (1961).

48. Huffaker, B. H. Experimental Studies on Predation: Dispersion Factors and Predator-Prey Oscillations. Hilgardia 27, 343–383 (1958).

49. Matsuo, T., Martínez‐Ramos, M., Bongers, F., Van Der Sande, M. T. & Poorter, L. Forest structure drives changes in light heterogeneity during tropical secondary forest succession. Journal of Ecology 109, 2871–2884 (2021).

